# A Benchmarking Study of Random Projections and Principal Components for Dimensionality Reduction Strategies in Single Cell Analysis

**DOI:** 10.1101/2025.02.04.636499

**Authors:** Mohamed Abdelnaby, Marmar R. Moussa

**Affiliations:** School of Computer Science University of Oklahoma, Norman, OK, USA

## Abstract

Principal Component Analysis (PCA) has long been a cornerstone in dimensionality reduction for high-dimensional data, including single-cell RNA sequencing (scRNA-seq). However, PCA’s performance typically degrades with increasing data size, can be sensitive to outliers, and assumes linearity. Recently, Random Projection (RP) methods have emerged as promising alternatives, addressing some of these limitations. This study systematically and comprehensively evaluates PCA and RP approaches, including Singular Value Decomposition (SVD) and randomized SVD, alongside Sparse and Gaussian Random Projection algorithms, with a focus on computational efficiency and downstream analysis effectiveness. We benchmark performance using multiple scRNA-seq datasets including labeled and unlabeled publicly available datasets. We apply Hierarchical Clustering and Spherical K-Means clustering algorithms to assess downstream clustering quality. For labeled datasets, clustering accuracy is measured using the Hungarian algorithm and Mutual Information. For unlabeled datasets, the Dunn Index and Gap Statistic capture cluster separation. Across both dataset types, the Within-Cluster Sum of Squares (WCSS) metric is used to assess variability. Additionally, locality preservation is examined, with RP outperforming PCA in several of the evaluated metrics. Our results demonstrate that RP not only surpasses PCA in computational speed but also rivals and, in some cases, exceeds PCA in preserving data variability and clustering quality. By providing a thorough benchmarking of PCA and RP methods, this work offers valuable insights into selecting optimal dimensionality reduction techniques, balancing computational performance, scalability, and the quality of downstream analyses.

## 1 Background and Motivation

Single-cell RNA sequencing (scRNA-seq) has revolutionized the field of genomics, enabling cellular and molecular profiling of gene expression at the individual cell level. This technology’s particular strength lies in uncovering cellular variability within samples and tissues, providing insights into dynamic biological processes, disease progression, and immune responses [1] [2]. However, scRNA-seq data, specifically the ‘count’ matrices are inherently high-dimensional and sparse, posing significant challenges for data analysis and interpretation [3].

Dimensionality reduction is a crucial step in scRNA-seq analysis that aims to transform high-dimensional data into a lower-dimensional latent space while preserving essential biological aspects. While distance-based methods, e.g. t-distributed stochastic neighbor embedding (tSNE)[4] or Uniform Manifold Approximation and Projection (UMAP)[5] focus on preserving locality (i.e. similarity between cells), other techniques have a strength in preserving or capturing variability; Principal Component Analysis (PCA) is one of the most widely used techniques for this purpose [6]. PCA reduces dimensionality by identifying linear combinations of variables (i.e. principal components) that capture the maximum variance of the data. It has been effectively applied in various scRNA-seq studies for visualization, clustering, and trajectory inference [7] [8][9].

Despite its widespread use, PCA has limitations when applied to large complex scRNA-seq data. PCA assumes strictly linear relationships among variables and struggles to capture non-linear structures inherent in biological data [10]. Additionally, PCA is sensitive to outliers and noise, which are common in scRNA-seq data due to technical variability and sparsity [11] [12], moreover, the PCA algorithm is typically computationally intensive for large sets [13].Random Projection (RP) methods have emerged as promising alternatives. Based on the Johnson-Lindenstrauss lemma [14], RP techniques reduce dimensionality by projecting data onto a lower dimensional subspace using a random matrix while aiming at approximately preserving pairwise distances [15]. While RP methods have shown success in fields like machine learning and signal processing [16], their application in scRNA-seq data analysis is only starting to gain attention.

Previous studies in scRNA-seq have primarily focused on comparing PCA with non-linear dimensionality reduction techniques like t-SNE and UMAP [17] [18]. Although PCA has been extensively benchmarked [19], these studies did not include RP methods, highlighting an opportunity for further investigation into the potential advantages of RP in scRNA-seq data analysis. Only a couple of recent studies have explored the use of RP in scRNA-seq, demonstrating its efficiency in handling large, complex datasets [20][21], yet, to our knowledge there are no systematic comprehensive benchmarking studies that contrast PCA and RP techniques. Such evaluation can shed the light on not only time or computing complexity of implementations, but also suitability for downstream analyses, hence the motivation for this work.

## 2 Approach

In this study, we systematically benchmark multiple PCA algorithms, including standard (or full) PCA and randomized SVD-based PCA, against two common RP methods: Sparse Random Projection (SRP) and Gaussian Random Projection (GRP). We assess their computational efficiency and effectiveness in downstream analysis tasks using both labeled (i.e with known ground truth of cell populations’ annotation) and unlabeled scRNA-seq datasets. By providing a comprehensive evaluation of PCA and RP methods, our work aims to:

- Benchmark the practical time complexity of PCA methods (full SVD and randomized SVD) and RP methods (Sparse and Gaussian) to determine their scalability on increasingly large scRNA-seq datasets.
- Investigate the effectiveness of each method in downstream analysis, specifically clustering.
- Evaluate each method’s ability to preserve data structure or locality as well as variability.

Clustering performance is evaluated using Hierarchical Clustering and Spherical K-Means algorithms. For labeled datasets, we measure clustering accuracy using the Hungarian algorithm and Mutual Information. For unlabeled datasets, we use the Dunn Index and Gap Statistic to assess cluster separation. We also examine the preservation of data variability using the Within-Cluster Sum of Squares (WCSS) metric.

***Our findings (see sections 4 and 5) demonstrate that RP methods not only offer significant computational speed-ups over PCA but also rival and, in some cases, surpass PCA in preserving latent structure, enhancing clustering performance. This study expands the toolbox of dimensionality reduction techniques available for scRNA-seq data analysis and underscores the importance of method selection and evaluation in the face of growing data complexity***.

## 3 Methods

As previously described, we evaluated two types of PCA: full SVD and randomized SVD and two common RP techniques were explored: Sparse Random Projection (SRP) and Gaussian Random Projection (GRP). SRP uses sparse random matrices, leading to faster computations and reduced memory usage [22]. GRP employs dense random matrices with entries drawn from a Gaussian distribution, providing theoretical guarantees on distance preservation. All methods were applied across varying target component sizes. For the commonly targeted range (a few, e.g. 5 to 25 components), we varied our tests in steps of 1, for the less explored range (25 to 1000), we varied our tests in steps of 25. We evaluated the practical implementation computational efficiency and clustering performance of these methods using labeled and unlabeled scRNA-seq datasets as described in 3.3. Below, we give a summary of PCA and RP definitions when applied to scRNA-Seq data:

### 3.1 PCA

#### Standard PCA

Standard PCA computes the principal components using the full SVD of the count matrix. Let **X** ∈ ℝ^*m*×*n*^ be the count matrix, where *m* is the number of observations or cells and *n* is the number of features or genes. The SVD of **X** is expressed as: **X** = **UΣV**^⊤^ where: **U** ∈ ℝ^*m*×*m*^ is an orthogonal matrix containing the left singular vectors, **Σ** ∈ ℝ^*m*×*n*^ is a diagonal matrix with singular values on the diagonal, and **V** ∈ ℝ^*n*×*n*^ is an orthogonal matrix containing the right singular vectors. The top *k* principal components are obtained by selecting the first *k* columns of **V**, denoted as **V**_*k*_. The data projected onto these principal components is given by: **T** = **XV**_*k*_. [23]

#### Randomized PCA

Randomized SVD-based PCA approximates the principal components efficiently by using randomization to reduce dimensionality followed by computing the SVD. This method leverages a random Gaussian matrix **Ω** ∈ ℝ^*n*×*k*^ to approximate the range of **X**: **Y** = **XΩ**, where **Y** ∈ ℝ^*m*×*k*^. An SVD is then computed on the smaller matrix 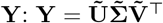. The approximate principal components are obtained from 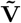, and the data projected onto these components is: 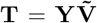. This approach significantly reduces computational complexity while claiming to maintain accuracy. [24]

### 3.2 RP

RP methods reduce the dimensionality of data by projecting it onto a random lower-dimensional subspace and is thought to be computationally efficient while claiming to preserve the structure of high-dimensional data. The two RP methods evaluated are:

#### Sparse RP

Sparse RP uses a sparse random matrix **R** ∈ ℝ^*n*×*k*^, where *k* is the target dimension. The entries of **R** are defined as:

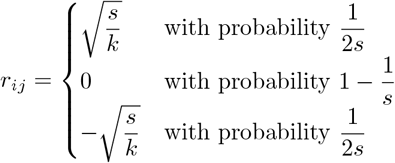

where *s* controls the sparsity of the matrix **R** (not to be confused with the count matrix sparsity). This scaling ensures that the expected value of 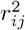 is 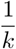, which is crucial for preserving distances during projection. The transformed data **Z** is then computed as: **Z** = **XR** Sparse RP methods are thought to be advantageous over GRP in computational efficiency and memory savings, especially when *s* is large. [22]

#### Gaussian RP

Gaussian RP uses{a den)se random matrix **G** ∈ ℝ^*n*×*k*^ with entries drawn independently from a Gaussian distribution: 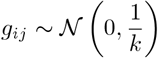 The projected data **Z** is obtained as: **Z** = **XG**[15] GRP is thought to be more effective in preserving the pairwise distances of the original data due to the properties of Gaussian distributions, which could translate into better performance in clustering tasks.

***Our study is set to explore these properties and claims of strength of each of the methods in practical scRNA-Seq data setting***.

### 3.3 Datasets

Four publicly available scRNA-seq datasets were used to evaluate the effectiveness of the dimensionality reduction methods. These include:

1. **Sorted PBMC Dataset** (from [25]): This dataset includes 2,882 cells and 7,174 genes and serves as a **labeled** set with 7 annotated distinct cell populations, providing a baseline for clustering methods.
2. **50/50 Mixture Dataset (Jurkat:293T Cell Mixture)** (from [26]): This dataset contains approximately 3,400 cells, with an approximately 50% distribution of Jurkat and 50% of 293T fairly homogeneous cell lines. This is a **labeled** dataset with two ground truth labels representing both cell lines.
3. **Targeted PBMC Dataset** (from [27]): This dataset utilizes a panel of putative immune-related genes (approximately 1000 genes after QC) and contains unannotated 10,497 cells (**unlabeled**). Beside for unbiased clustering, this dataset was additionally used with varying sizes ranging from 1000 to 10,000 cells to evaluate scalability.
4. **COVID-19 T Cell Dataset** (from [28]): This data focuses on human TCells in the context of bronchoalveolar immune cell from COVID-19 patients and healthy subjects and is an unlabeled dataset.

The **Sorted PBMC Dataset, 50/50 Mixture Dataset (Jurkat:293T Cell Mixture)**, and **COVID-19 T Cell Dataset (Liao 2020)** were all downloaded from the SC1 Tool. The **Targeted PBMC Dataset** was downloaded from the 10x Genomics website.

### 3.4 Validation Metrics

As described in 2. Clustering performance is evaluated using two clustering methods to evaluate PCA and RP robustness when controlling for the effect of the clustering algorithm:

- **Agglomerative Hierarchical Clustering** was used with Ward linkage algorithm and Cosine distance, constructing dendrograms or trees that were cut to the known number of clusters in case of labeled sets.
- **Spherical K-Means Clustering** was used also with Cosine distance, grouping cells based on directional similarities.

To test accuracy and effectiveness or quality of the clustering analysis as one of the main downstream analyses for scRNA-seq data, we measured the following metrics:

- **Accuracy (Hungarian algorithm)**- used for known Ground Truths, maximizing the sum of label matches between predicted and true labels.
- **Mutual Information**- used for known Ground Truths, given as 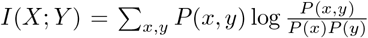, where *P* (*x, y*) is joint probability of a true label *x* and predicted *y*. Higher values indicate better match.
- **Dunn Index**- used for unknown Ground Truths and easures the ratio between inter-cluster distance and intra-cluster compactness. It is given as 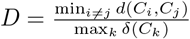, where *d*(*C*_*i*_, *C*_*j*_) is the distance between cluster centers *C*_*i*_ and *C*_*j*_, and *δ*(*C*_*k*_) is the diameter of cluster *C*_*k*_. Higher values mean better separation.
- **Gap Statistics**- also used for unknown Ground Truths and is given by 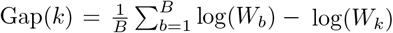, where *W*_*k*_ is the within-cluster dispersion for *k* clusters, and *W*_*b*_ represents the expected dispersion under a null reference distribution. Here too, higher values indicate better separation.

We also examine the preservation of data variability using the Within-Cluster Sum of Squares (WCSS) defined as 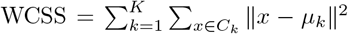, where *K* is the number of clusters, *C*_*k*_ is the set of cells in cluster *k, x* is a data point in *C*_*k*_, and *μ*_*k*_ is the centroid of cluster *k*.

## 4 Results

### 4.1 Dimensionality Reduction for Downstream Analysis

We examined two aspects of the dimensionality reduction properties of the examined methods; first how they are used for visualizing the data and secoond and more importantly, how well cells cluster in the reduced low dimensional space:

#### Visualization

Figure 1 visualizes the Sorted PBMC dataset using PCA and GRP using only the first two components each. Although both methods show overlap in the projections, GRP provides less clear visualization compared to PCA, this is expected since with the PCA the first few components capture more of the data latent properties like variability while this distinction between the earlier and later components is meaningless for RP methods.

**Fig. 1.**
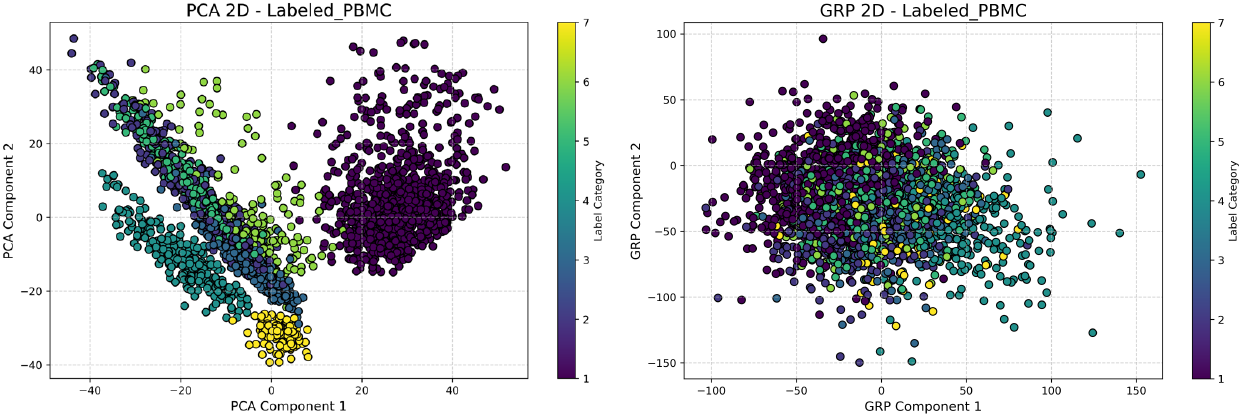
Visualization for the Sorted (color codes used for labels) PBMC dataset using PCA (left) with Full SVD and GRP (right), both using the first two components only.

#### Evaluating Clustering Accuracy and Quality

We used clustering effectiveness as means of evaluating how well the reduced, low dimensional latent space produced from different methods is suited for use in downstream analyses. Figure 2 shows the results for the labeled sets displaying the Dunn Index and Mutual Information for all labeled sets and all methods over varying number of components.

**Fig. 2.**
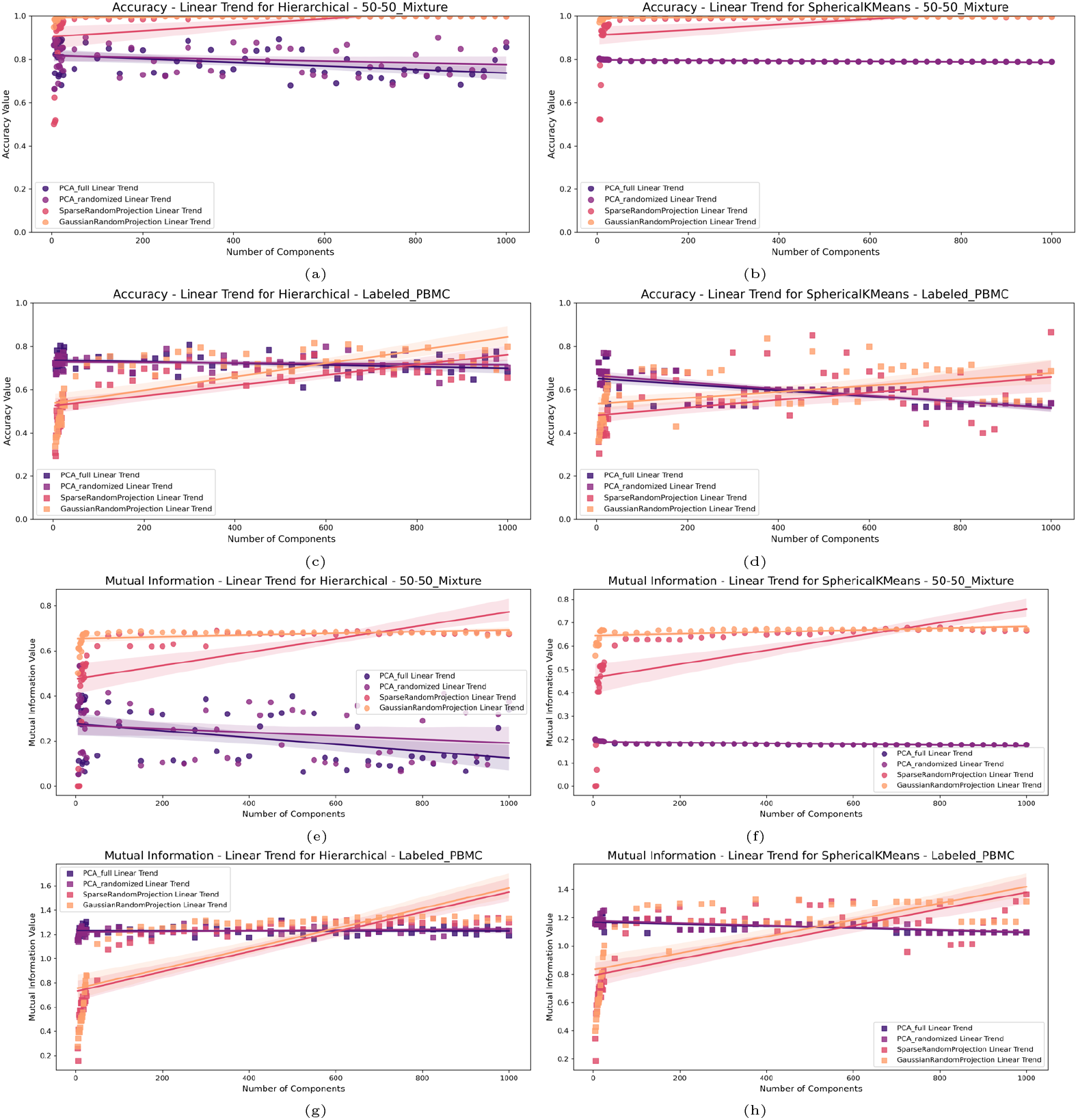
a) to d) : Accuracy for a) Jurkat - 293T 50-50 Mixture dataset with Hierarchical Clustering b) Jurkat - 293T 50-50 Mixture with SKMeans c) Labeled PBMC with Hierarchical Clustering d) Labeled PBMC with SKMeans. e) to h): Mutual Information for e) Jurkat - 293T 50-50 Mixture with Hierarchical Clustering f) Jurkat - 293T 50-50 Mixture with SKMeans g) Labeled PBMC with Hierarchical Clustering h) Labeled PBMC with SKMeans.

Furthermore, for unlabeled data, we examined the ‘goodness’ of clustering by evaluating the Dunn Index and Gap Statistics values as described in Methods. Figure 3 illustrates these results, again for all evaluated methods and varying number of components used for clustering.

**Fig. 3.**
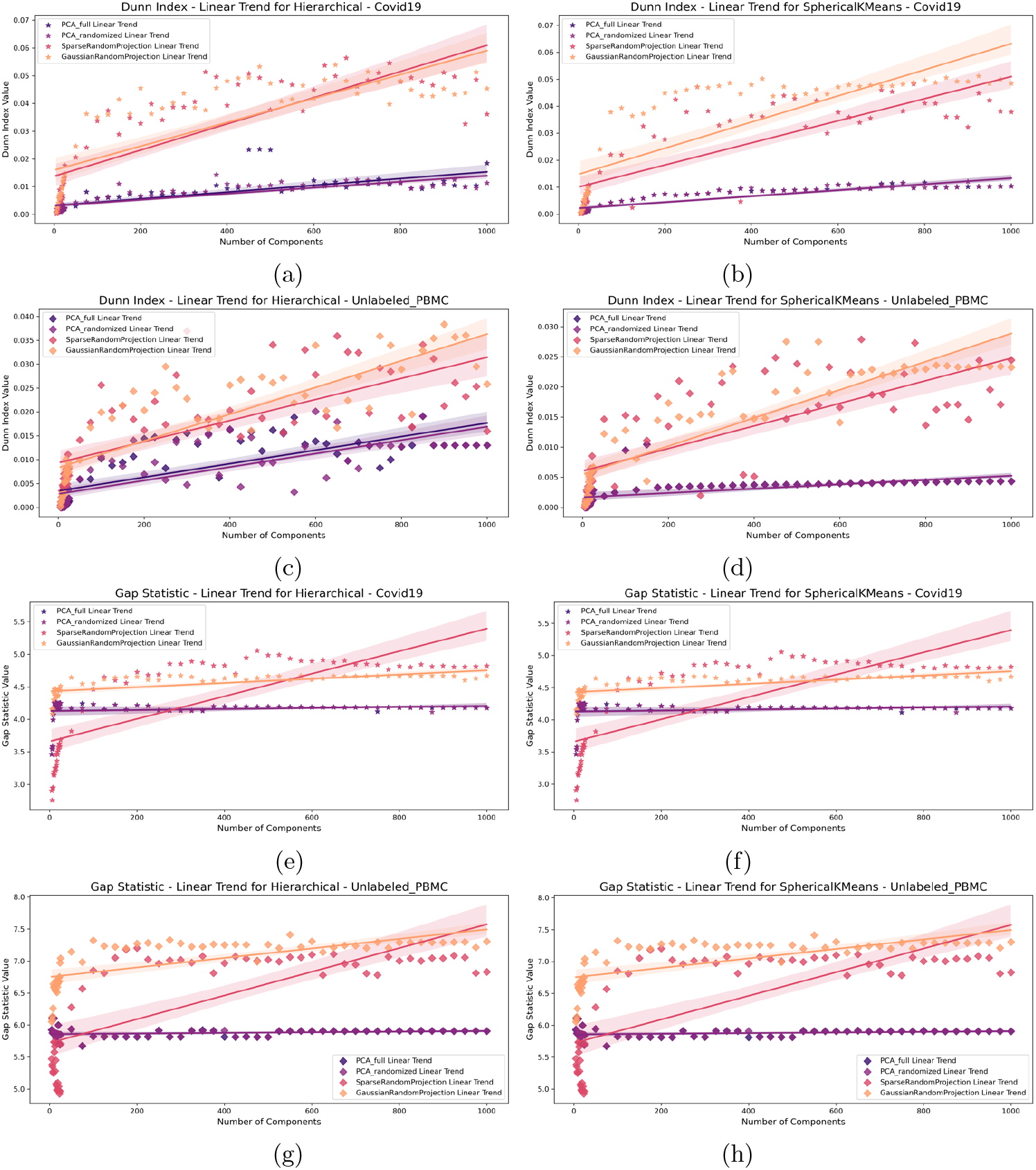
a) to d) : Dunn Index for a) Covid19 dataset with Hierarchical Clustering b) Covid19 dataset with SKMeans c) Unlabeled PBMC dataset with Hierarchical Clustering d) Unlabeled PBMC dataset with SKMeans. e) to h): Gap Statistic for e) Covid19 dataset with Hierarchical Clustering f) Covid19 dataset with SKMeans g) Unlabeled PBMC dataset with Hierarchical Clustering h) Unlabeled PBMC dataset with SKMeans.

### 4.2 Variability Preservation

Since capturing variability or heterogeneity is one of the main insights PCA and RP methods provide, we set to evaluate whether preserving variability would directly be correlated with perhaps lower accuracy performance, however when evaluating the WCSS metric to measure the heterogeneity ‘preserved’ by each dimensionality reduction method for each dataset, see Figure 4, we see that RP methods indicate higher variability preservation while still achieving higher accuracies, especially when using *>* 25 components for clustering.

**Fig. 4.**
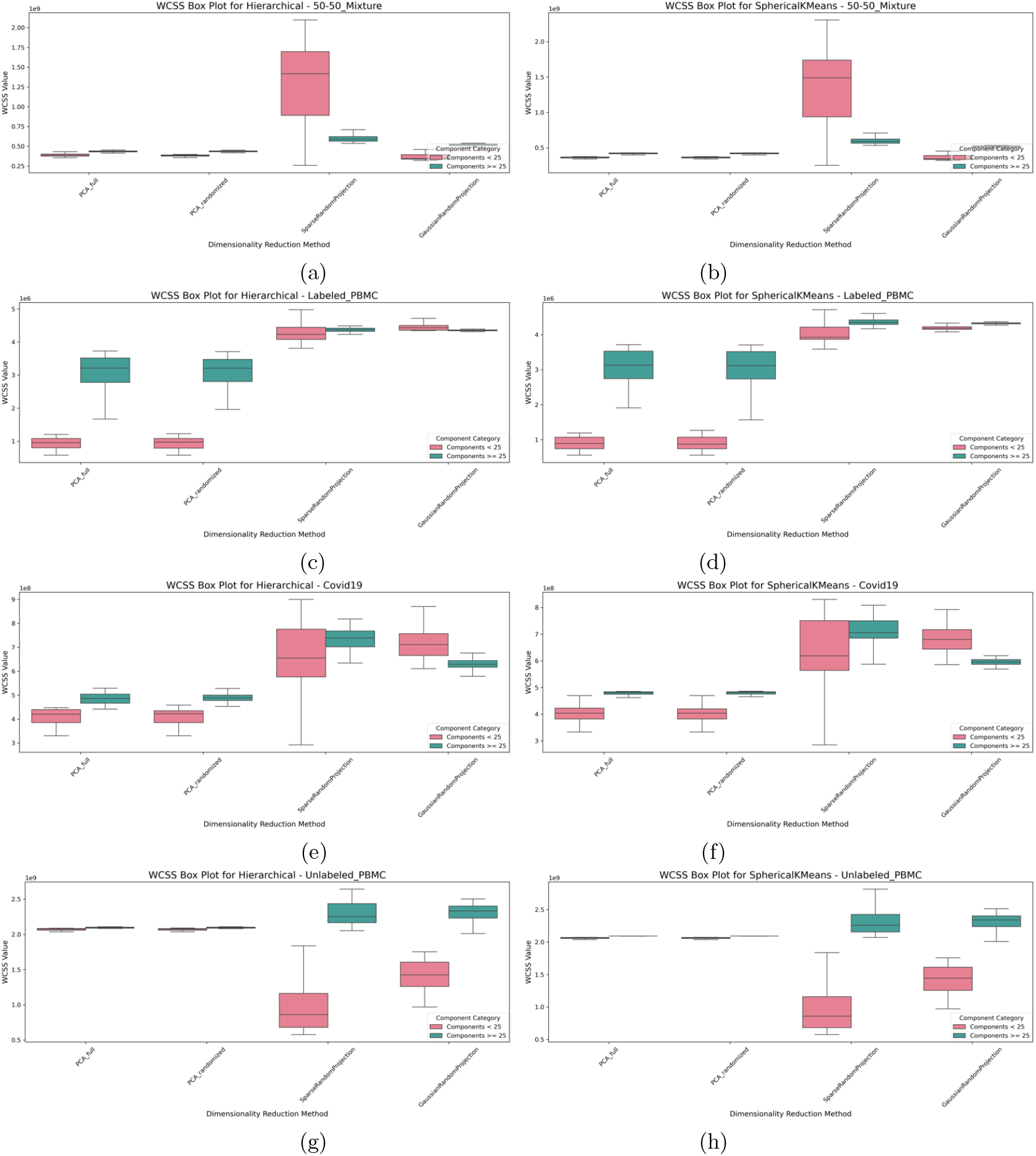
a) to d): WCSS (Variability Measure) for a) Jurkat - 293T 50-50 Mixture with Hierarchical Clustering b) Jurkat - 293T 50-50 Mixture with SKMeans c) Labeled PBMC with Hierarchical Clustering d) Labeled PBMC with SKMeans. e) to h): WCSS (Variability Measure) for e) Covid19 with Hierarchical Clustering f) Covid19 with SKMeans g) Unlabeled PBMC with Hierarchical Clustering h) Unlabeled PBMC with SKMeans.

### 4.3 Locality Preservation

We compared the ability of PCA and RP to preserve locality or pairwise similarity of cells by projecting the Sorted PBMC dataset and the Jurkat - 293T 50-50 Mixture dataset into a 2D space using UMAP.

Figure 5 shows that SRP results in visually similar UMAP to PCA, preserving essential data relationships crucial for downstream analyses, such as clustering or other. To quantify the locality preservation further, we calculated cluster accuracy metrics for PCA and RP when using 500 embeddings or components each followed by applying UMAP to project further to a 3 dimensions UMAP space. SKMeans Clustering accuracy and Mutual Information metrics using the resulting UMAP components are given in table 1.

**Table 1.** Clustering Performance Metrics at 500 Components and after UMAP to 3D.

| Method | Metrics at 500 Components |  |
| --- | --- | --- |
|  | Accuracy | Mutual Information |
| PCA Full | 0.7893 | 0.1784 |
| PCA Randomized | 0.7893 | 0.1784 |
| GRP | 0.9970 | 0.6713 |
| SRP | 0.9921 | 0.6499 |
| Method | Metrics After UMAP to 3D |  |
|  | Accuracy | Mutual Information |
| PCA Full + UMAP | 0.9967 | 0.6707 |
| PCA Randomized + UMAP | 0.9949 | 0.6593 |
| GRP + UMAP | 0.9878 | 0.6329 |
| SRP + UMAP | 0.9973 | 0.6739 |

**Fig. 5.**
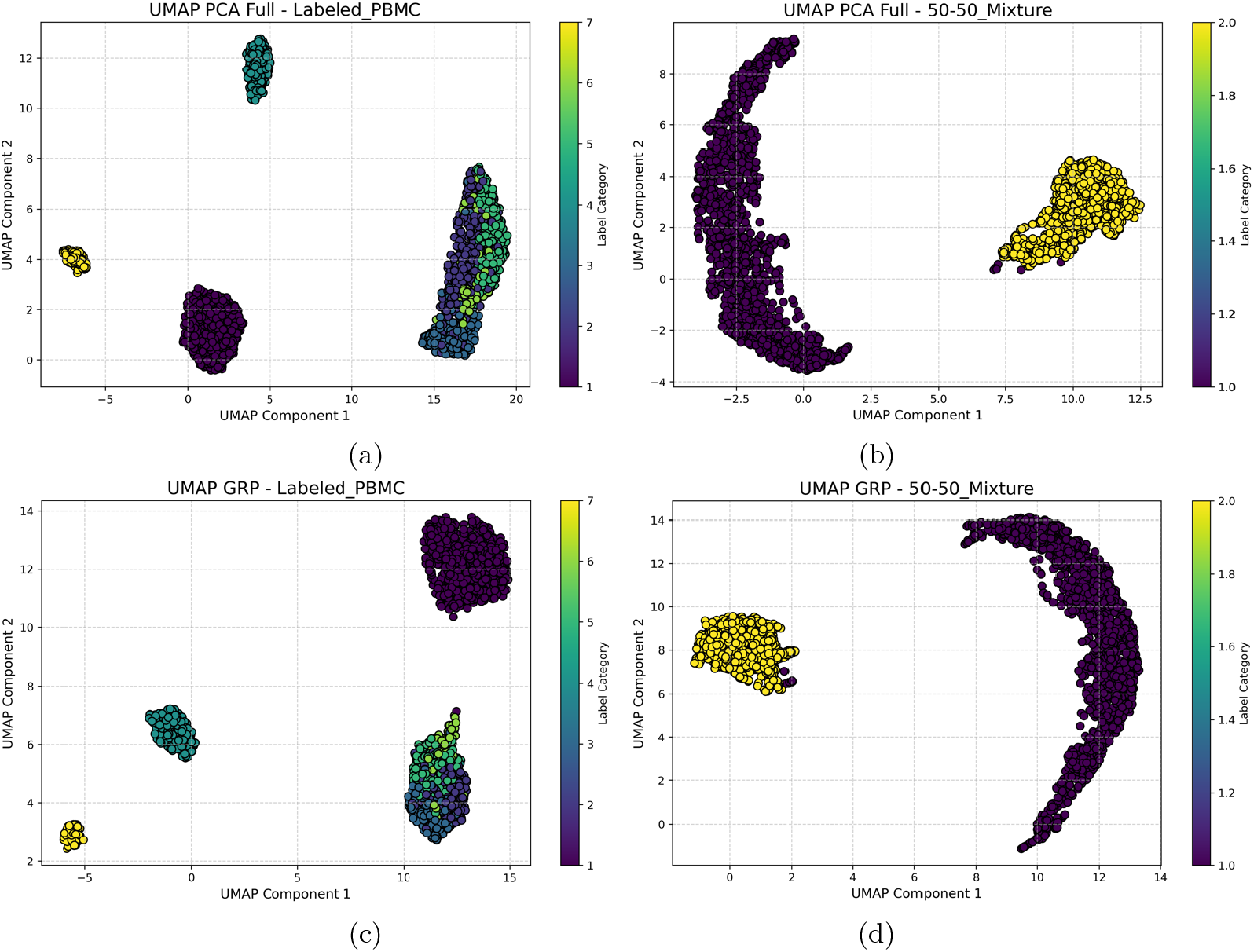
UMAP 2D projection of the Sorted PBMC dataset with dimensionality reduction of PCA and GRP of 500 components. a) PCA with full SVD for the Labeled PBMC dataset b) PCA with full SVD for the Jurkat - 293T 50-50 Mixture dataset c) GRP for the Labeled PBMC dataset d) GRP for the Jurkat - 293T 50-50 Mixture dataset

### 4.4 Computational Efficiency

Figure 6 (a to d) demonstrate the execution times of the dimensionality reduction methods across all the datasets. Additionally, we conducted an experiment in which we varied the size of the targeted PBMCs by Gibbs sub-sampling cells to create multiple datasets of varying sizes ranging from 1000 to 10,000 cells with steps of 1000; using these datasets we assessed the execution time in relevance to dataset size. Figure 6 (e) highlights the scalability of RP methods across increasing dataset sizes, with SRP consistently being the most efficient method, even for largest sample sizes.

**Fig. 6.**
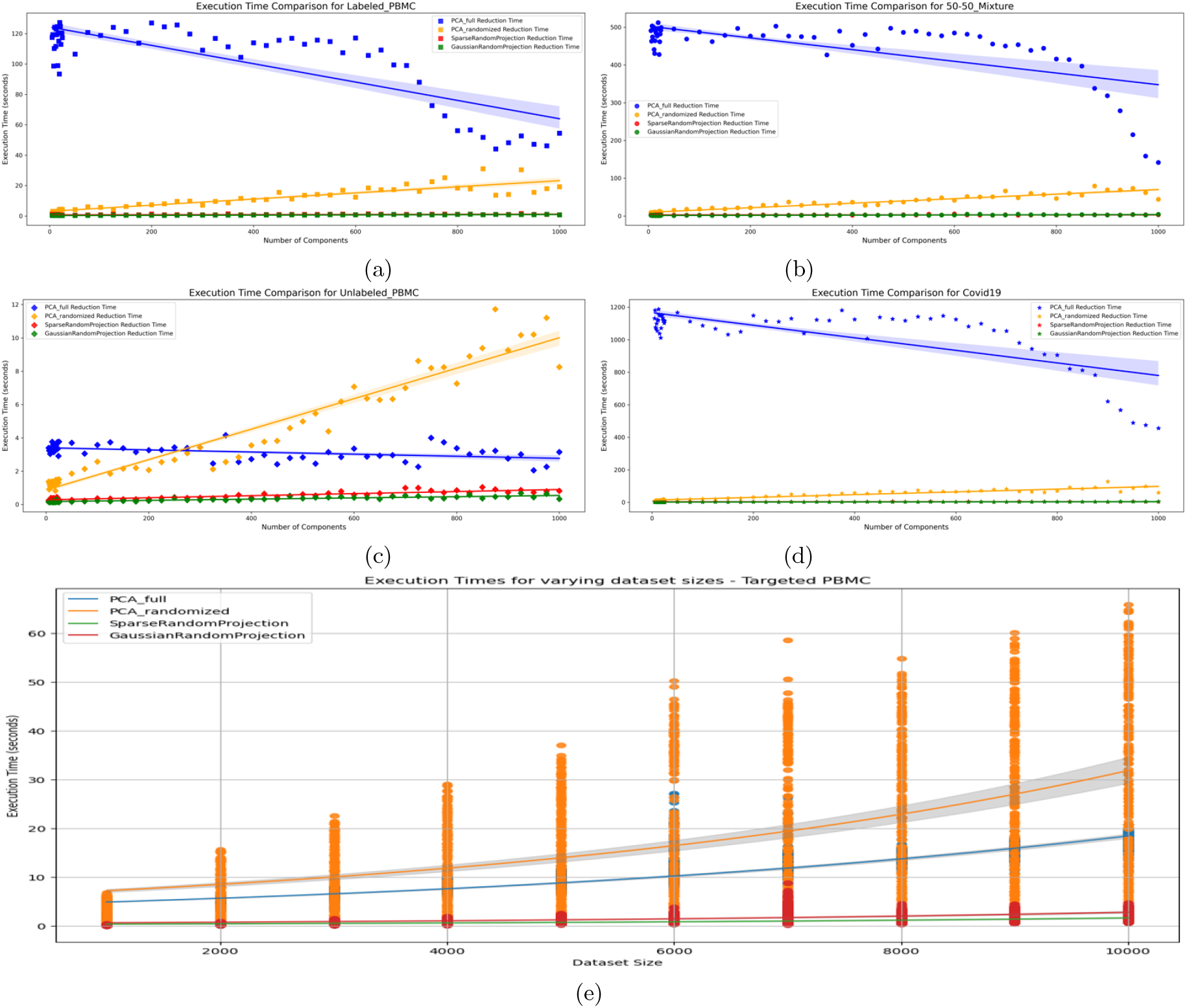
Execution time vs. number of components for each dimensionality reduction technique on the a) Sorted PBMC dataset, b) Jurkat - 293T 50-50 Mixture, c) Targeted PBMC dataset, and d) Covid19 data. In e) we show Execution time vs. dataset sizes (1000 to 10,000 samples) for each of the dimensionality reduction techniques.

## 5 Discussion

Our findings indicate that RP outperforms PCA in clustering accuracy across several datasets. As shown in Figure 2, both SRP and GRP variants of RP achieved higher accuracy compared to PCA (utilizing Full and Randomized SVD) on the Jurkat-293T 50-50 Mixture dataset, using both Hierarchical and SKMeans clustering methods. In the labeled PBMC dataset, PCA performed slightly better in lower-dimensional spaces, but as dimensionality increased, RP either matched (for Hierarchical clustering) or outperformed PCA (for SKMeans).

This trend is similarly reflected in when considering he Mutual Information Index, suggesting RP’s superiority in the 50-50 Mixture dataset. While RP initially under-performs in lower dimensions in the labeled PBMC dataset, it begins to surpass PCA as more components are added, suggesting RP’s increasing reliability with higher dimensions.

For the unlabeled datasets, RP again demonstrates improved performance over PCA. Figure 3 shows that RP consistently outperformed PCA in Dunn index values on the Covid19 dataset across both clustering algorithms. In the unlabeled PBMC dataset, RP and PCA had nearly identical performance in lower dimensions, but RP gained an advantage as dimensionality increased for both clustering algorithms.

In Figure 3, we evaluated clustering separation with the Gap Statistic on the unlabeled datasets, where RP again consistently outperforms PCA. These results support the effectiveness of RP, particularly in higher dimensions and across diverse datasets, underscoring its potential as a superior dimensionality reduction method for clustering tasks compared to PCA.

In Figure 4, the box plots reveal that RP methods generally display higher WCSS variability compared to PCA, especially in the range using higher number of components (*>* 25) (Note the median line in all box plots). This increased WCSS median value reflects a broader spread within the clusters formed by RP, which, while resulting in slightly looser clusters, does not detract from RP’s overall clustering accuracy. Interestingly, as dimensionality increases, the variability of the WCSS measure itself (inter-quartile range of the box plots) decreases for RP, suggesting that RP’s clustering performance becomes more stable and consistent at higher dimensions, still preserving more variability than PCA (higher median line).

Indeed, for all labeled data as well as for the Covid19 dataset, the box plots further show RP’s tendency toward higher WCSS across dimensions. Despite this variability, RP consistently performs well in clustering accuracy, as supported by other evaluation metrics. For the Unlabeled PBMC dataset, a targeted gene panel data set, both PCA and RP show relatively similar WCSS values in higher dimensions, but RP’s variability is decreased increases when only a few components are used, reflecting the importance of all genes in the panel for capturing the latent variability of this dataset.

Finally, Figure 6 illustrates RP’s clear and significant advantage in terms of execution time. We measured the execution time across all datasets for different numbers of components; RP, especially GRP, consistently and significantly outperformed both PCA methods, highlighting the strength of this method from the computational complexity perspective. We noticed an interesting phenomenon, where PCA with Full SVD shows some decrease in execution time at higher dimensions. This can be explained by the effect of sparsity in the produced embeddings with higher components (i.e. when higher dimensions are calculated, the per component mean value is lower as shown in Figure 7. Another interesting observation, in (c), the lower dimension targeted panel dataset, the randomized PCA performs better than Full SVD PCA up to a certain point before becoming less efficient, highlighting the overhead of performing randomization whwn not needed for lower dimensional data.

**Fig. 7.**
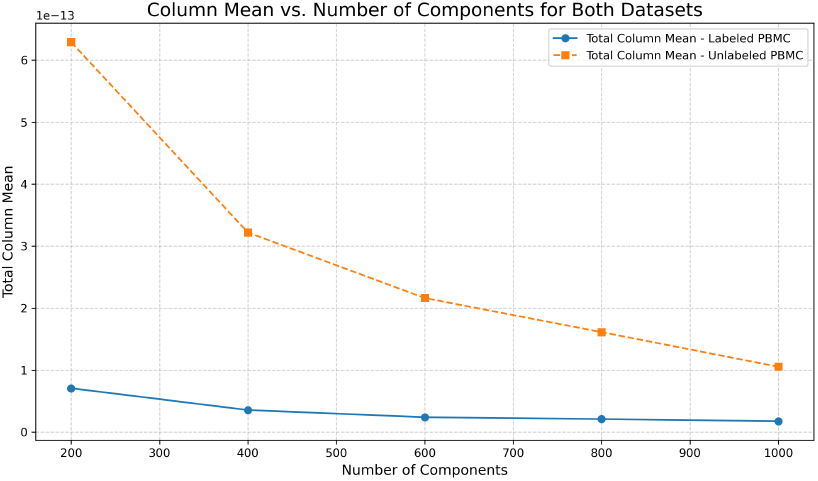
Mean value per component vs. Number of Components calculated for Labeled and Unlabeled PBMC Datasets, highlighting the impact of PCA Full SVD on execution time.

All in all, and across all datasets, RP demonstrates superior performance in execution time and downstream analysis, highlighting its practical advantages and value in single cell RNA-Seq analysis.

## 6 Conclusion

This study demonstrates that RP can outperform PCA in scRNA-Seq analysis, especially, clustering tasks across various datasets, particularly as more dimensions are considered. RP, especially the SRP and GRP variants, achieved higher clustering accuracy and faster execution times compared to PCA. Although PCA performed slightly better in lower dimensions on some datasets, RP consistently excelled in higher-dimensional spaces, showing strong results across accuracy metrics like the Mutual Information Index and Dunn Index. While RP sometimes resulted in broader cluster spreads, this variability did not compromise clustering performance, particularly in high-dimensional settings.

## Data and Code Availability

The data and code used for this analysis are available on GitHub: https://github.com/moussa-lab/BenchmarkingPCA-RP or upon reasonable request to the authors.

## Acknowledgments

This work is supported by the following awards: NSF-2341725, NIH-NCI K25CA270079, and OU-BIC2.0.

